# Prom1 expression does not mark a stem/progenitor population in the mouse oviduct epithelium

**DOI:** 10.1101/2020.08.19.257923

**Authors:** Matthew J Ford, Yojiro Yamanaka

## Abstract

The oviduct or fallopian tube is the site of fertilization and preimplantation embryonic development. The epithelium lining the oviduct consists of multiciliated and secretory cells, which support fertilization and preimplantation development, however, its homeostasis still remains poorly understood. CD133/*Prom1* has been used to identify adult stem cell populations in various organs and often associated with cancer stem cell property. Using a Cre-recombinase based lineage tracing strategy, we found that CD133/*Prom1* expression was not associated with a stem/progenitor population in the oviduct but marked a sub population of multiciliated and secretory cells which did not propagate. Interestingly, *Prom1* expressing secretory cells rapidly transition to multiciliated cells and progressively migrate to the tips of epithelial folds in the ampulla. Our results show that CD133/*Prom1* expression cannot be used as a progenitor/stem cell marker in the mouse oviduct.

## Introduction

*Prom1* (Prominin-1) is a transmembrane glycoprotein whose expression was first characterized on the surface of hematopoietic stem cells using the antigen CD133 (1,2). Prom1/CD133 is now recognized as an adult stem cell marker and has be used to isolate various tissue-specific stem cell populations across multiple organs (3–5). CD133 has also been identified on the surface of cancer cells with stem-like properties in many tumors including ovarian, liver, brain, prostate, colon, hepatocellular and lung cancer (6). The specificity of Prom1/CD133 as an adult stem cell marker has however come into question due to conflicting reports of CD133 specification, potentially due to the use of different antibodies between studies, and the broader expression pattern seen in *Prom1* reporter mouse lines, indicating potential differences between CD133 antibody stainings and *Prom1* expression (7–13). In a recent study Zhu and colleagues performed a lineage tracing experiment of *Prom1* expressing cells and found variation in the generative and proliferative capacity of *Prom1* expressing cells between organs (13). It was also found that the *Prom1* expressing cells with a high generative capacity had a higher risk of tumor formation after an oncogenic insult, indicating that the stem-cell characteristics of *Prom1* expressing cells varies between different organs and that locating those populations with a high generative capacity may identify highly susceptible cells to malignant transformation.

The oviduct epithelium is constituted of multiciliated and secretory cells, that aid the transport, function and survival of gametes and embryos (14). It has been shown by lineage tracing studies that secretory cells are proliferative and can differentiate into multiciliated cells, suggesting that secretory cells may represent a bipotent progenitor (15). Mounting evidence suggests that these cells are also the cell-of-origin in many cases of high-grade serous ovarian carcinoma (HGSOC), the most common and lethal form of ovarian cancer (16). The presence of an adult stem cell, ie, an undifferentiated cell whose progeny replenish dying cells, has not be characterised in the oviduct. Multiple studies have however eluded to the presence of a resident adult stem by the identification of label-retaining cells and organoid forming cells, enriched in the distal oviduct (17–20). The expression of several common adult stem cell markers have also been reported such as *Cd44*, *Prom1* and *Lgr5* (21–23). However, there have been no *in vivo* characterisations of these populations that confirm their status as a progenitor or adult stem cell.

In this study we investigate the stem cell properties of *Prom1* expressing cells in the mouse oviduct using lineage tracing techniques. We find that *Prom1* expression is limited to a sub population of multiciliated and secretory cells along the length of the oviduct. These cells were not proliferative and did not propagate over time. *Prom1* expressing secretory cells rapidly differentiated to multiciliated cells and progressively moved from the base to tip of epithelial folds over time in the ampulla but remain restricted to the base of epithelial folds in the isthmus. Taken together our results show that *Prom1* expression does not mark a resident adult stem cell population in the mouse oviduct.

## Results

### CD133/Prom1 becomes restricted to a sub population of multiciliated and secretory cells in the mouse oviduct during development

In order to determine the expression pattern of *Prom1* in mouse oviduct epithelial cells we used an antibody targeted to the CD133 antigen. Prior to epithelial differentiation we detected CD133 staining on the apical surface of all epithelial cells at E19.5 (embryonic day 19.5) (Figure 1A). In adult mice CD133 was restricted at the apical surface of a subset of PAX8 positive and PAX8 negative cells sparsely distributed along epithelial folds of the distal oviduct (infundibulum and ampulla) (Figure 1B). In the isthmus, staining was concentrated to the apical surface of PAX8 positive epithelial cells at the base of epithelial folds (Figure 1B). CD133 was also broadly expressed in organoids grown from isolated mouse oviduct epithelial cells, localising to the apical surface of multiciliated and secretory cells (Figure 1C). To estimate the proportion and identify which cell types express *Prom1* we used Prom1^C-L^:Rosa26^Tdtomato^ mice (12,24). Five doses of tamoxifen were administered on consecutive days followed by a 72-hour lag before oviducts were dissected. In the distal oviduct 7.64 ±2.35% (n = 3 mice) of epithelial cells were Tdtomato positive and sporadically distributed throughout the epithelium (Figure 1D). This was significantly higher than the proportion labeled with CD133 (2.85±1.8% n = 3 mice), suggesting the CD133 antibody could be detecting a specific isoform or posttranslational modification. In the isthmus, 10.37 ±4.5% (n = 3 mice) epithelial cells were labeled and located in clusters at the base of epithelial folds. No labeled cells were detected in the uterotubal junction (Figure 1D). In all regions Tdtomato was expressed predominantly in multiciliated cells and did not show colocalization with proliferation marker Ki67 (0 ki67+/Tdtomato+ cells, n = 3 mice) (Figure 1E).

**Figure 1:**
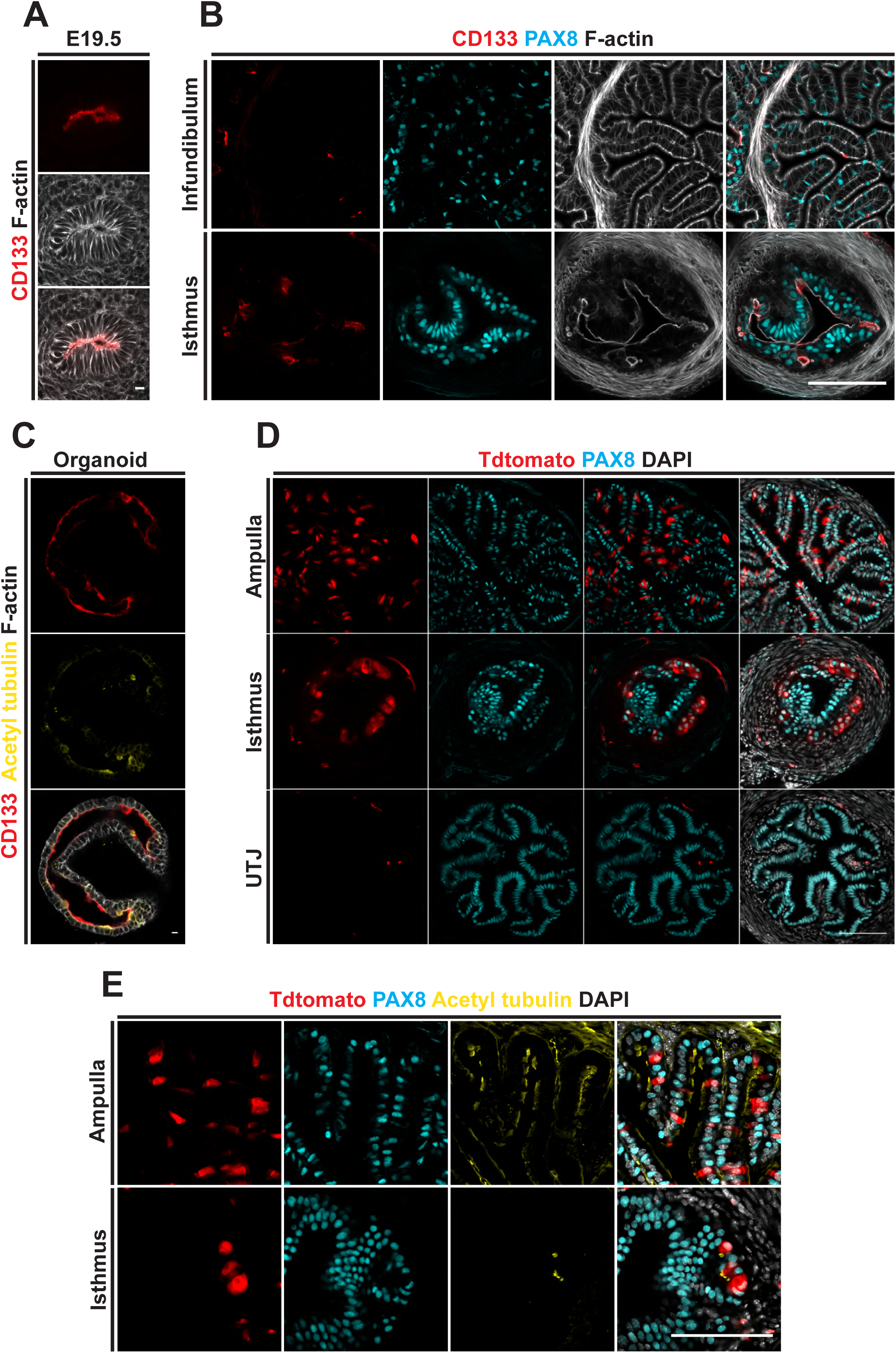
Distribution of CD133/*Prom1* expressing cells in the mouse oviduct. **(A)** A CD133 antibody was used to locate *Prom1* expressing cells in the mouse oviduct. CD133 staining was detected at the apical surface of all epithelial cells on E19.5 prior to epithelial cell differentiation. **(B)** In the adult oviduct CD133 was found on the apical surface of sparsely distributed PAX8+ and PAX8- cells in the distal oviduct but restricted to the apical surface of cells at the base or epithelial folds in the isthmus. **(C)** Broad staining of CD133 was found on multiciliated and secretory cells in differentiated mouse oviduct organoids. **(D)** The distribution pattern of Tdtomato+ cells in Prom1^C-L^:Rosa26^Tdtomato^ mice 72 hours after 5 injections of tamoxifen. No Tdtomato+ cells were detected in the uterotubal junction. **(E)** The majority of Tdtomato+ cells were multiciliated in the distal oviduct and isthmus. Scale bar in A and C = 10µm, B, D and E = 100µm.

### *Prom1* labels a subset of multiciliated cells and differentiating secretory cells with a low generative capacity

To investigate the stem cell properties of *Prom1* expressing cells we used a lineage tracing approach with Prom1^C-L^:Rosa26^Tdtomato^ mice. Five injections of tamoxifen were administered over consecutive days and oviducts collected over a 5-month period (Figure 2A). We detected no significant change in the proportion of Tdtomato positive cells in the ampulla and a slight decrease in labeled cells in the isthmus (n = 3 mice per time point) (Figure 2B–D). To follow the differentiation of *Prom1* expressing cells over time we calculated the intensity of PAX8 expression, a secretory cell marker, in labeled cells. In the ampulla 20% of cells had a visually detectable level of PAX8 staining after 72 hours and were distributed in all regions of epithelial folds but proportionally enriched at the base (Figure 2E). Of these PAX8 positive cells the majority had a relativity low level of PAX8 staining compared to the total epithelial cell population (Figure 2F and G). Over the course of the lineage tracing experiment we detected a rapid decrease in the proportion of PAX8 expressing Tdtomato+ cells to 1% after 1 month (Figure 2F). We were unable to confirm this observation in the isthmus however due to the ubiquitous expression of PAX8 in all epithelial cells.

**Figure 2:**
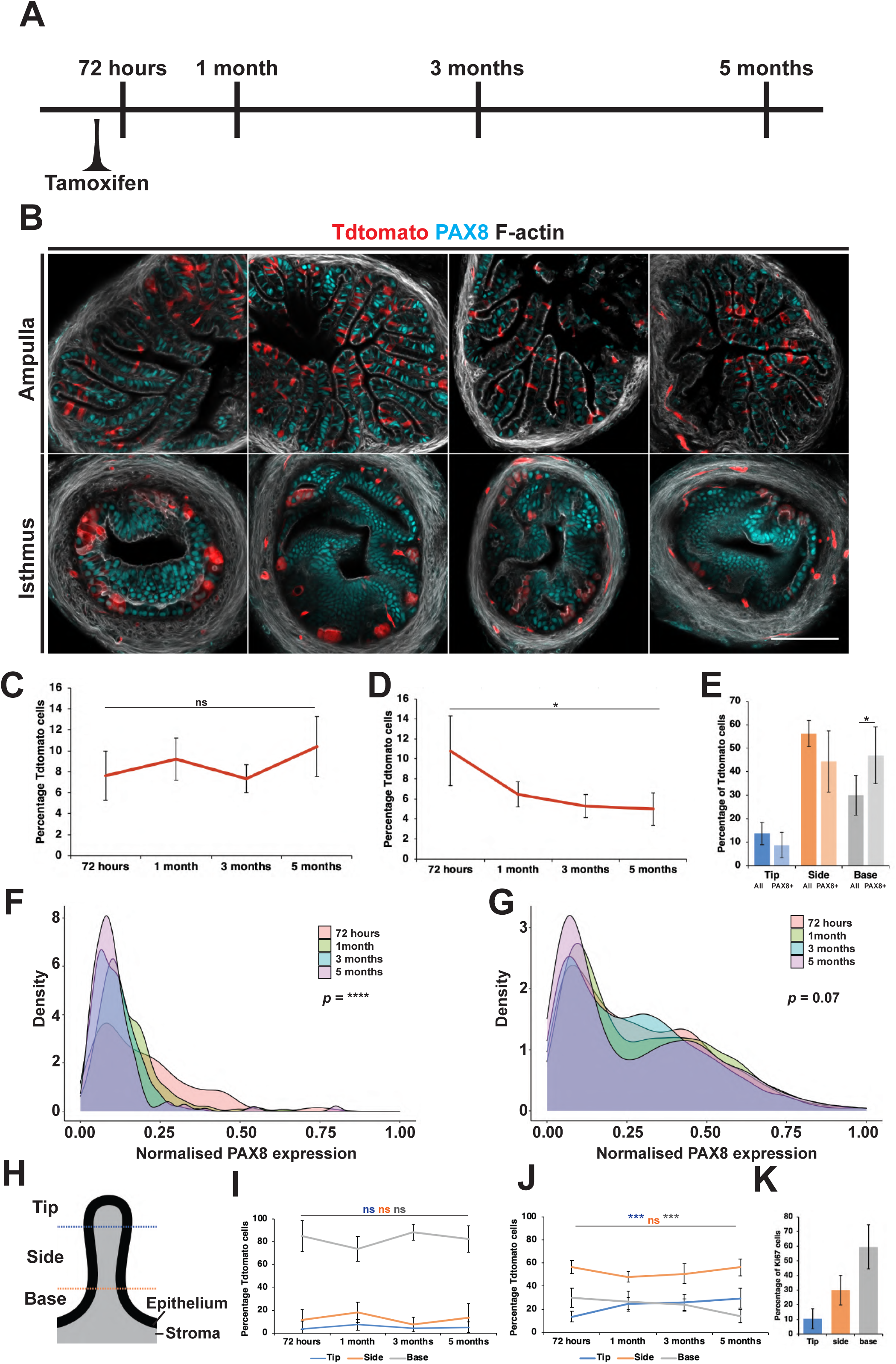
Lineage tracing *Prom1* expressing cells in the mouse oviduct. **(A)** Lineage tracing was performed in Prom1^C-L^:Rosa26^Tdtomato^ mice by administration of 5 doses of tamoxifen on consecutive days followed by collection of oviduct at different time points over a 5-month period. **(B)** Selected examples showing the distribution pattern of labeled cells over 5-months in the ampulla and isthmus. **(C)** No significant change in the proportion of Tdtomato+ cells was detected in the ampulla. **(D)** In the isthmus a steady drop in the proportion of Tdtomato+ cells was seen. **(E)** In the ampulla Tdtomto+ cells were found in all regions of epithelial folds. Tdtomato+ cells that also had visible PAX8 staining were also found in all regions but were overrepresented at the base of epithelial folds. **(F)** Density plot showing the distribution of PAX8 expression in Tdtomato+ cells over 5-months lineage tracing. Low PAX8 expression was detected in a subpopulation of cells at 72 hours. After 1-month almost all Tdtomato cells had no detectable levels of PAX8 expression (n = 238 cells per time point). **(G)** Density plot showing the distribution of PAX8 expression of all epithelial cells from the same lineage experiment (n = 2438 cells per time point). **(H)** Schematic of an oviduct epithelial fold in a transverse section showing the segmentation method used to classify epithelial cell position. **(I)** Line graph showing the proportion of Tdtomato+ cells within each region throughout the lineage tracing in the isthmus, showing no significant changes. **(J)** In the ampulla we detected a significant increase in the proportion of Tdtomato+ cells at the tip and a decrease at the base over 5-months. **(K)** Quantification of the proportion of Ki67+ cells by location, showing the majority to be located at the base of epithelial folds (n = 5 mice, 226 cells). Scale bar in B = 100µm. Statistical tests used in C-E,I and J = two-tail students T-test, F and G = two-sample Kolmogorov-Smirnov test comparing 72 hour and 1 month distributions. N = 3 mice for all lineage tracing data.

### Multiciliated cells migrate to the tip of epithelial folds in the ampulla

As *Prom1* expressing cells were not proliferative we were able to track the movement of labeled cells in the mouse oviduct to investigate the processes of homeostasis in the ampulla and isthmus. The position of Tdtomato positive cells was determined in relation to the base, side and tip of epithelial folds in transverse sections (Figure 2H). In the isthmus labeled cells remained restricted to clusters of cells at the base of epithelial folds (Figure 2I). In the ampulla however, we detected a progressive increase in the proportion of labelled cells at the tip of epithelial folds and a decrease at the base (Figure 2J). Suggesting a base to tip drift of epithelial cells over time. In keeping with this observation, the majority of proliferating cells (Ki67+) in the ampulla were detected away from epithelial tips (Figure 2K).

## Conclusion

CD133/Prom1 is a well characterised adult stem and cancer stem cell marker in many tissues (1–6). The utility of CD133/Prom1 as a general adult stem cell across tissues has however come into question with reports of broader Prom1 expression in transgenic mouse lines and variations in generative capacity of *Prom1* expressing cells across tissues (7–13). In the presented study we investigated the stem cell characteristics of *Prom1* expressing cells in the mouse oviduct. We determined that *Prom1* expression becomes restricted to a sub population of multiciliated cells and secretory cells during development. These cells were not proliferative and were dispersedly distributed in the distal oviduct while being restricted to clusters of cells at the base of epithelial folds in the isthmus, in concordance with the distribution pattern of multiciliated cells. Using lineage tracing we showed that the *Prom1* expressing population did not expand over time and in the case of the isthmus decreased over the course of the experiment. Taken together our results show that *Prom1* does not label a stem or progenitor population in the mouse oviduct.

Multiciliated cell turnover rate in the distal mouse oviduct has been estimated to be around 6 months (25). As we see a near exclusive labeling of multiciliated cells one month after tamoxifen injections we would have expected to see a steady drop in the number of labeled cells over time. However, in the ampulla we identified no significant change in the proportion of labeled cells over 5 months. We also found that *Prom1* labels a population of cells in the ampulla with low PAX8 expression that rapidly transition into multiciliated cells. Considering the slow turnover of *Prom1* expressing cell it is possible that *Prom1* may mark PAX8+ cells differentiating into multiciliated cells but is subsequently lost from aged multiciliated cells. The differences between the ampulla and isthmus may reflect a broader expression of *Prom1* in the isthmus multiciliated population or a faster turnover rate.

The oviduct is morphologically segmented into four distinct regions containing varying proportions of multiciliated and secretory cells (26). The homeostasis of the entire oviduct epithelium is generally considered as a single population. Several reports have however started to highlight differences between the distal and proximal epithelial cell populations, showing a concentration of label retaining cells, cells expressing known adult stem cell markers and organoid forming cells in the distal region of the oviduct (17,18,20,22,23,27). In our lineage tracing study, we find that the distribution pattern of *Prom1* expressing cells and the drift of these cells is distinct between the ampulla and isthmus. This result supports the notion of independent homeostatic mechanisms within the morphologically distinct regions of the oviduct.

## Methods

### Mouse stains and injections

All animal work was approved by the internal ethics committee at McGill University and undertaken at the Goodman Cancer Research Centre animal facility. *Prom1C-L* (#017743) and *Tdtomato*^flox/flox^ mice (Ai14, #007914) were acquired from JAX and maintained on a mixed C57BL/6, CD1 background. C_57_BL/6 stock mice were used as wild type mice. Tamoxifen was prepared on the day of injection at a concentration of 20 mg/ml in corn oil and 4 mg per 25 g administered by intraperitoneal injection.

### Immunofluorescence

Embryonic Müllerian and adult oviducts were dissected in PBS and fixed in 4% w/v PFA/PBS (Polysciences) for 30 minutes at room temperature followed by 3 PBS washes. Ducts were cryoprotected through a sucrose/PBS gradient 4°C, allowing time for the ducts to sink to the bottom of the tube between each gradient. Ducts were embedded in OCT mounting solution (Fisher HealthCare) and snap frozen on dry ice before being stored in a −80°C freezer. 10µm sections were cut using the microtome cryostat Microm HM525 (Thermofisher) mounted on Superfrost glass slides, air dried and stored in −80°C freezer. For immunofluorescence, sections were allowed to thaw for 10mins at room temperature followed by rehydration with PBS. Sections were permeabilized for 5 minutes with 0.5% v/v Triton-X/PBS (Sigma) and then blocked with 10% v/v Fetal Bovine Serum (FBS) (Wisen Bioproducts), 0.1% v/v Triton-X in PBS for 1 hour at room temperature. Primary antibodies were diluted in 1% v/v FBS, 0.1% v/v Triton-X in PBS to working concentrations and incubated overnight in a dark humidified chambered at 4°C. The primary antibodies used in this study were: PAX8 (proteintech #10336-1-AP), acetylated tubulin (Sigma Aldrich #T7451) and CD133 (ThermoFisher, monoclonal 13A4 #14-1331-80). The followed day sections were washed before incubation with secondary antibodies (ThermoFisher, Alexa Fluor, diluted 1/400 in 1% v/v FBS, 0.1% v/v Triton-X in PBS) and for 1 hour in a dark humidified chambered at room temperature. In some cases, Phalloidin (ThermoFisher, 1/500) was added to label f-actin. Sections were then washed again with 5µg/ml DAPI (Fisher Scientific) added to the final wash step before sections were mounted with prolong gold (Invitrogen) for imaging. For lineage tra
cing, fixed samples were embedded in 4% agarose sectioned using a vibratome into 100µm sections. The sections were then permeabilized for 10 minutes in 0.5% v/v Triton-X/PBS, stained as above and then mounted in prolong gold between two coverslips separated by a spacer (Invitrogen Sercure-seal 0.12mm).

### Mouse oviduct epithelial cell organoid culture

Mouse oviduct epithelial cells were isolated by Fluorescence-activated cell sorting (FACS). Oviducts were disassociated into single cells by incubation in collagenase B (5mg/ml) and DNase I (5U/100ul) in DMEM (Gibco) containing 100 IU/ml of Penicillin and 100ug/ml of Streptomycin for 35 minutes at 37°C followed by passing through a needle series. The resulting single cell suspensions were passed through a 40µm cell strainer, centrifuged in a table top centrifuge at 1,500rpm for 5 minutes and resuspended in 200µl staining solution containing conjugated antibodies Ep-CAM-APC (#118213 Biolegend) and Brilliant violet 785-CD45 (#103149 Biolegend) diluted 1/150 with DMEM (Gibco) containing 100 IU/ml of Penicillin and 100ug/ml of Streptomycin and 1% FBS for 15 minutes in the dark on ice. Cell suspensions were then centrifuged, washed in PBS and resuspended in FACS media (DMEM (Gibco) containing 100 IU/ml of Penicillin and 100ug/ml of Streptomycin and 20% FBS). FACS was performed with the FACSAria Fusion (BD Biosciences) using FACSDIVA (Version 8). Cell suspensions were spiked with 1µl 5mg/µl DAPI (Thermofisher) just prior to sorting to identify dead or dying cells. Live Epcam+/CD45-epithelial cells were sorted into FACS media and washed in PBS before being seeded into a drop of Geltrex (Thermofisher). The Geltrex was allowed to set for 40 minutes in a 37°C humidified incubator with 5% CO_2_ before being submerged in IntestiCult media (STEMCELL technologies).

### Imaging and statistics

All imaging was performed on an LSM 800 confocal microscope (Zeiss) at the advanced bioimaging facility (McGill University). Image analysis was conducted with FIJI using custom designed software (28). Statistical analysis was then performed either using Excel (Microsoft) or R-studio.

## Declaration of interests

The authors declare no competing interests.

## Notes

### Competing Interest Statement

The authors have declared no competing interest.

